# Exploring Urban Street Green Perception from the Perspective of Combining GVI and NDVI: A Case Study of Zhongshan City, Guangdong Province

**DOI:** 10.1101/2023.05.21.541659

**Authors:** Lei Su, Weifeng Chen, Yan Zhou, Lei Fan, Junying Li

## Abstract

Urban street greening has a positive impact on the health of citizens and the urban environment. This study takes the representative streets in the main urban area of Zhongshan City, Guangdong Province as an example to explore urban street greening perception from the perspective of combining Green visual index (GVI) and Normalized difference vegetation index (NDVI). This study uses a deep learning based image semantic segmentation method to analyze Baidu Street View to calculate the GVI of the street, and uses GF-1 satellite data to calculate NDVI to compare and analyze the characteristics and correlation of GVI and NDVI of urban streets. The results show that: 1. The GVI of streets in the central urban area of Zhongshan varies from 8.06% to 36.00%, with Xingzhong Road in Shiqi District Street having the highest GVI; 2. The mean value of NDVI of each street shows different changes with the increase of buffer scale, and the mean value of NDVI has a strong scale sensitivity; 3. The highest Pearson correlation coefficient between GVI and 25m DNVI mean value was 0.862; 4. The GVI prediction model based on NDVI is: y=0.8249x+0.0181, R^2^=0.7433. On this basis, the shortcomings of street landscape are analyzed and optimization suggestions are given, providing reference for urban street landscape evaluation, spatial optimization, and landscape improvement.

## Introduction

Urban street greening has a positive impact on the health of citizens and the urban environment (Puppala et al. 2022). Street greening can not only alleviate the physical and mental pressure of residents, regulate the urban microclimate (Bowler et al. 2010), improve air quality (Janhäll 2015), but also alleviate the urban heat island effect (Gago et al. 2013), reduce noise, and control runoff caused by precipitation (Li et al. 2015). The public’s perception of the urban street greening significantly affects the public’s mental health (Van den Bosch and Sang 2017). Therefore, researchers have begun to pay attention to the relationship between street greening quality, green perception, and its influencing factors (Wu et al. 2020; Chen et al. 2019; Zhu et al. 2022). As one of the important urban spatial indicators, the Green visual index (GVI) reflects the amount of street greening, helping to reveal the level of street vegetation from a human perspective (Sun et al. 2023). GVI refers to the proportion of green plants within a person’s visual field. It was first proposed by Yoji Aoki (1987) to measure the greening construction of three-dimensional urban space. With the development of panoramic photography, Web street view images have become the main data source in the study of GVI (Lu 2019). Google Street View (GSV) has become the most popular data warehouse for scientific researchers (Griew et al. 2013), but because Google does not provide street view maps in Chinese Mainland at this stage, Baidu Street View (BSV) and Tencent Street View (TSV) have become alternative products in China. Yang et al. used Python to call Baidu Street View’s API in the form of HTTP URL to obtain street view photos in four directions, and calculated the GVI in the main urban areas of Fujian based on the Python image processing library OpenCV (Yang et al. 2022). Wan and Dai (2021) directly used the street view data capture method to study typical roads in the old urban area of Nanjing City. They used Matlab to convert the street view map into HSV color space to extract photo channel values, and extract the green part to calculate the GVI. This method is more accurate than manual processing. Wang et al. (2022) used OpenCV to stitch the GSV images of Savannah into 360° panoramic images, and used artificial intelligence semantic segmentation tool SegNet to calculate the Panoramic green view index, to explore the spatial distribution law of PGVI and analyze the influencing factors of street space, such as vegetation type, vegetation quantity and density, and street slope.

Although Web street view images have created convenience for the study of urban street landscape, there are some limitations in the use of street view map data(Smith et al. 2021; Machicao et al. 2022), such as: it is not guaranteed that street view images can be obtained in every street in every period. Therefore, other data with geographical coordinates, such as remote sensing index, are needed to supplement the data set. In the past, it was very common to obtain green vegetation index from remote sensing data (Xue and Su 2017), such as NDVI (Normalized difference vegetation index), which is the most commonly used vegetation remote sensing index. At present, some scholars have paid attention to the relationship between GVI and NDVI. For example, Li et al. (2021) used a grid of 250 m×250 m as the basic unit to study the spatial distribution difference between GVI and NDVI in urban streets; Lv et al. (2022) used Landsat data set as data source, added 50m to both sides of each road as buffer to calculate the average NDVI of each road, and then analyzed the relationship and influencing factors between GVI and NDVI. Larkin and Hystad (2019) downloaded 254 panoramic images of Google Street View from

Portland, Oregon, while studying the streetscape exposure assessment method for visible green spaces. They set the GSV dimension/longitude coordinate range of 50 meters, and calculated the ratio of GVI to NDVI to express the vertical dimension of green space. The above studies are all based on the statistics of NDVI within a fixed scale grid or buffer, ignoring the scale effect of NDVI as a landscape remote sensing index. In order to scientifically study the relationship between GVI and NDVI, it is necessary to set up more detailed spatial scale changes. High resolution remote sensing images will be more conducive to studying the correlation between NDVI and GVI at different scales.

This study takes the central urban area of Zhongshan as an example, using BSV images, the 91 satellite image software, and a graphical semantic segmentation method based on deep learning to obtain green visual index GVI, and obtaining a normalized vegetation index NDVI based on GF-1 remote sensing images. Further, the correlation between GVI and NDVI mean value at different scales is studied, and on this basis, the shortcomings of street landscape are analyzed and optimization suggestions are given.

## Materials and Methods

### Study area

Zhongshan City is located in the lower reaches of the Pearl River Delta, bordering Guangzhou in the north, Hong Kong and Macao, and facing Shenzhen across the sea. It is located in the geometric center of Guangdong-Hong Kong-Macao Greater Bay Area, with a land area of 1783.7 km^2^ (Figure 1). Zhongshan is in the south of the Tropic of Cancer, on the northern edge of the tropics, with abundant sunshine, abundant heat, warm climate and evergreen seasons. The main urban area of Zhongshan includes East District Street, West District Street, South District Street, Shiqi District Street, Port Town, Torch Development District Street and Wuguishan Street. In this study, the representative streets of the main urban area are selected as the research area. This area is the most densely populated, and the street view data is relatively perfect.

**Figure. 1.**
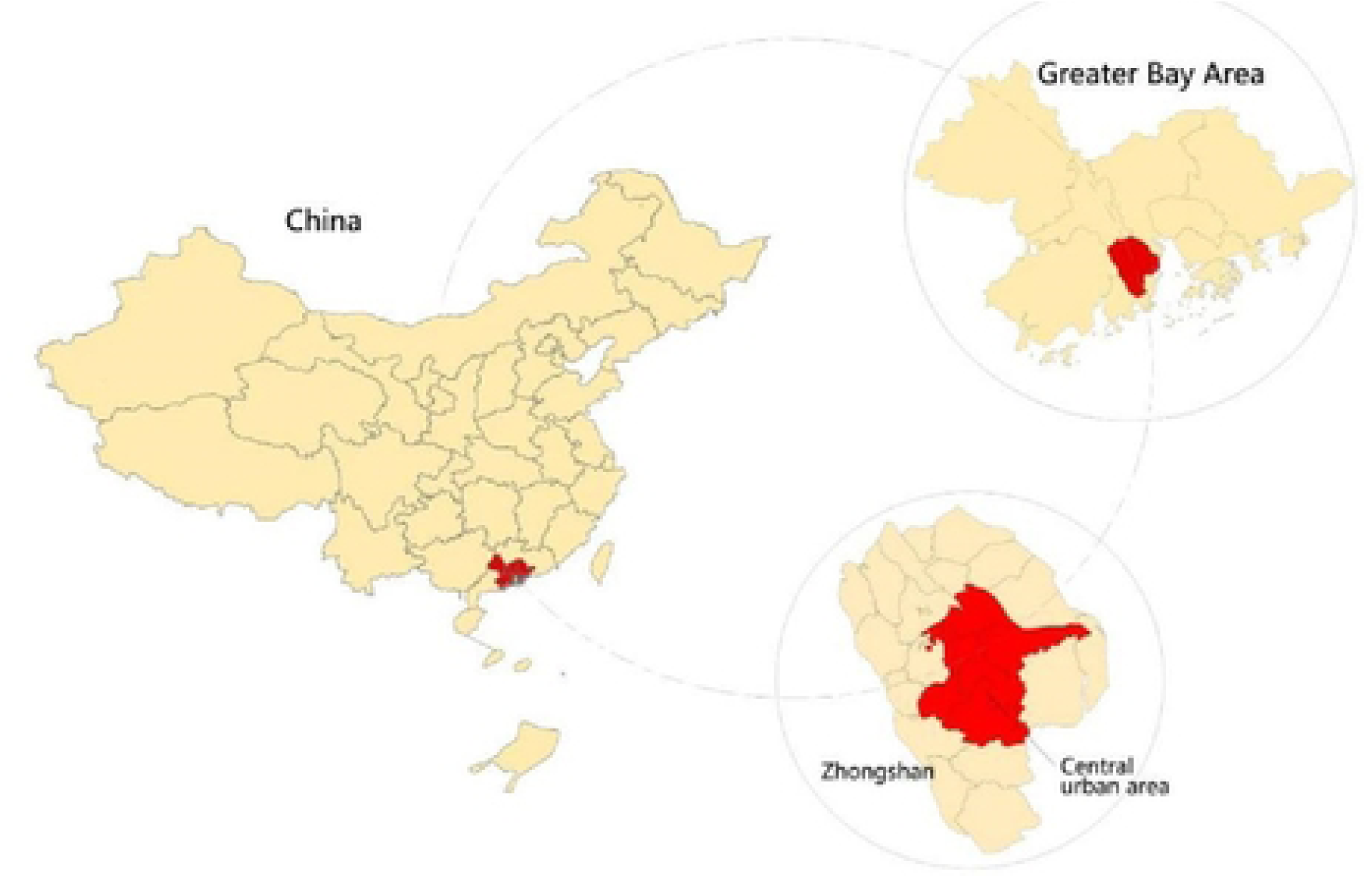
Study area.

### Observation point setting and data collection

Fully considering the characteristics of the current situation of land use and observation points, such as representativeness, universality, and operability, a representative street is selected from each District Street (Town) in the central urban area of Zhongshan as the research object (Figure 2), which are the main streets in urban administrative and commercial centers, national development zones, ecological reserves, major industrial platforms, and transportation hubs. They are representative of the District Street (Town) to which they belong and have distinctive planning characteristics, and are suitable as models for this study. The specific research objects include: Kiu Wan Road of East District Street, Chengnan 1st Road of South District Street, Choi Hung Avenue of West District Street, Xingzhong Road of Shiqi District Street, Hongle Avenue of Torch Development Zone Street, Chenggui Highway of Wuguishan District Street, Port Avenue of Port Town.

**Figure. 2.**
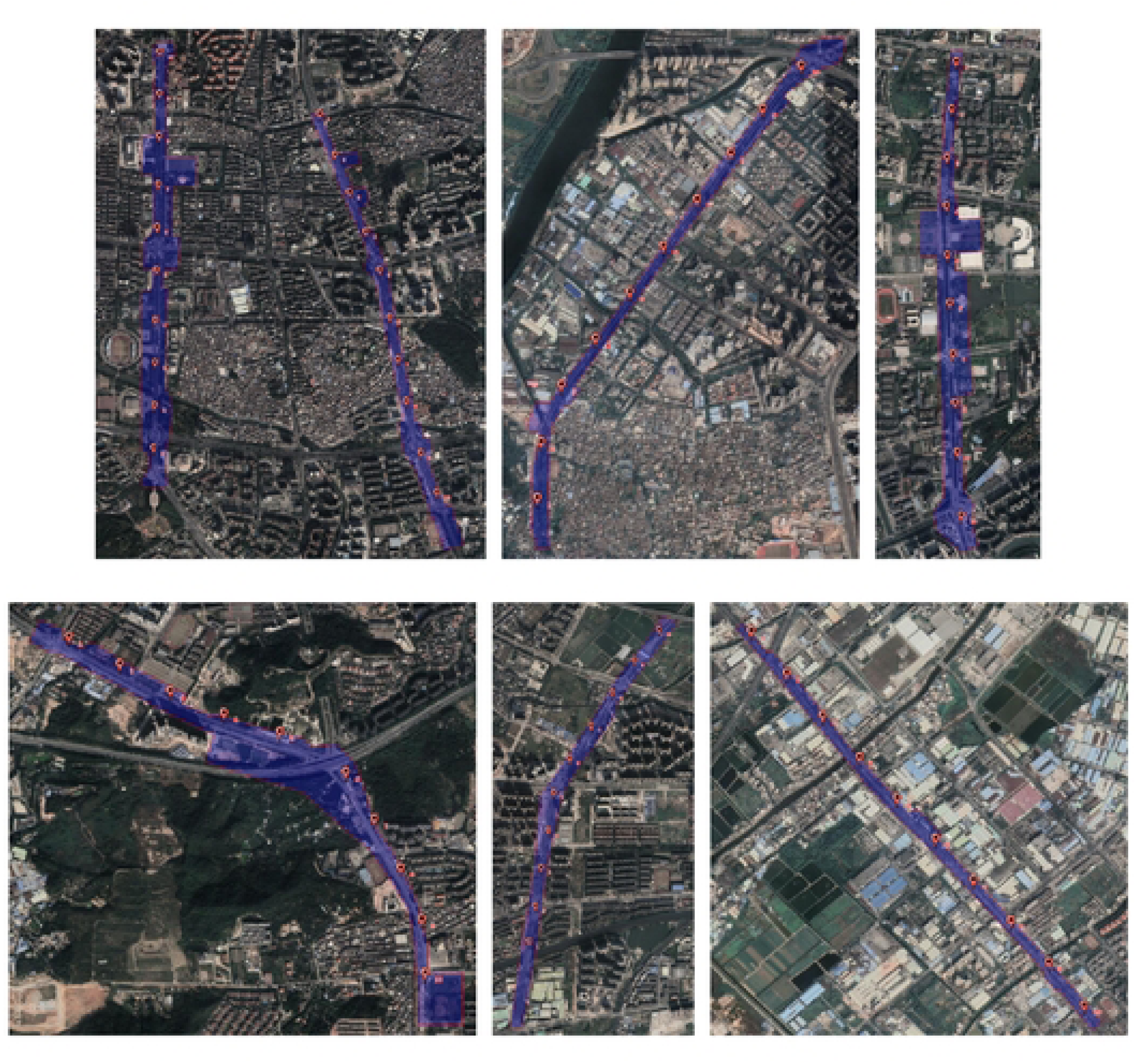
Satellite images of the study object and distribution of observation points.

It is more practical to reflect people’s feelings about urban street greening through street view pictures, and it is more convenient to obtain data by using street view pictures, which is not affected by many uncontrollable factors. These are effective means to measure the built environment. In this study, the street data in the central urban area of Zhongshan were obtained by using 91 Satellite Map Enterprise Edition, and the observation points were set up by using Arcgis software (10 observation points were set in each street), and the distance between observation points was 250m (the farthest distance for human eyes to see the outline of objects was 250m).

According to the coordinates of observation points, python software was used to collect street view photos in BSV map in four directions of 0°, 90°, 180° and 270° respectively (Figure 3), with a uniform size of 480×320px, and each photo has information such as the coordinate position and horizontal angle of the corresponding sampling point. A total of 70 coordinate points were selected to obtain 280 street view pictures.

**Figure. 3.**
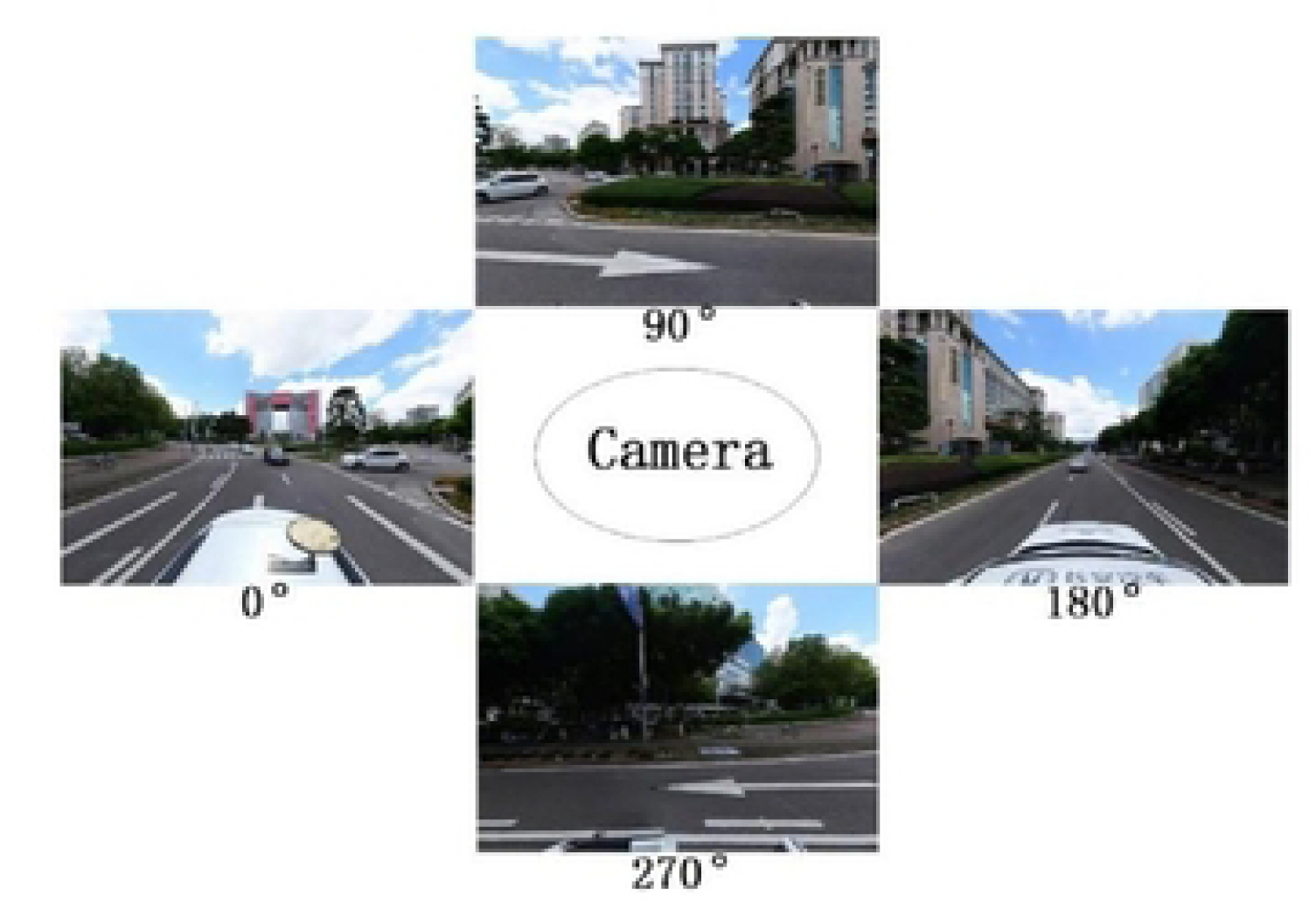
Example of green view index observation images.

### Image semantic segmentation method based on deep learning

Image semantic segmentation (ISS) is an interdisciplinary subject involving computer vision, pattern recognition, artificial intelligence and other research fields. Its definition is that each pixel in an image is assigned a pre-defined label representing its semantic category (Perronnin 2011), which can get the information that the image itself needs to express according to its texture, scene and other high-level semantic features, and has more practical value. This study adopts the image semantic segmentation method based on pixel classification, which belongs to the fully supervised learning image semantic segmentation method, specifically the image semantic segmentation method based on deep Fully Convolutional Network (FCN). FCN was proposed by Jonathan Long et al. of UC Berkeley in paper “Fully Convolutional Networks for semantic segmentation” in 2015 (Long et al. 2015). FCN is mainly used for image segmentation, to solve the problem that it is difficult for Convolutional Neural Network (CNN) to achieve fine (pixel-level) segmentation in image feature extraction. The frame structure of FCN is shown in Figure 4. FCN can transform the network originally used for image classification into the network used for image segmentation (Tian et al. 2019).

**Figure. 4.**
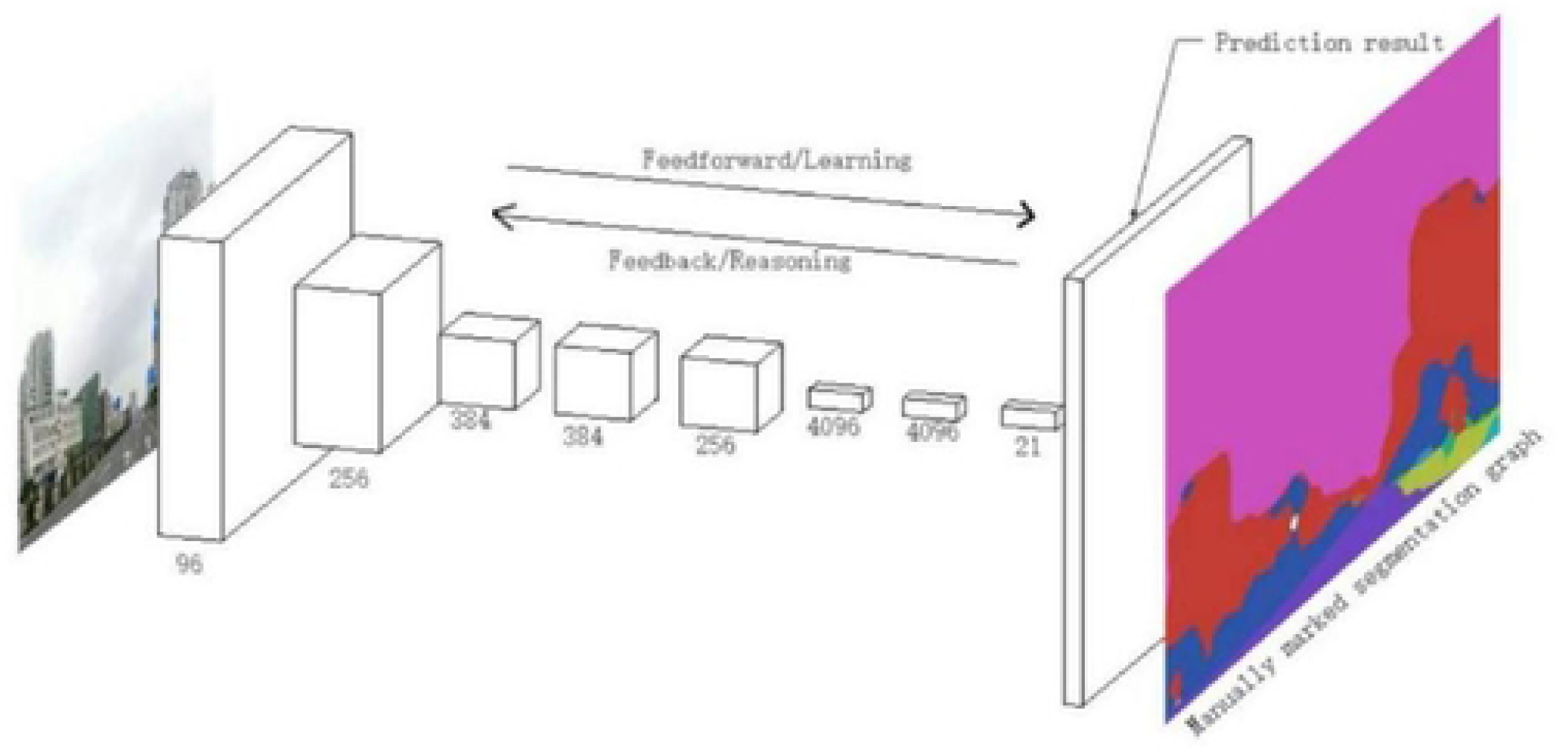
FCN frame structure.

### Extraction of green plants and calculation of GVI

In this paper, the deep learning FCN visual image semantic segmentation software (http://guihuayun.com/) based on ADE_20K data set training is selected to extract green plants. The software has a huge training set, accurate identification and stable operation, and can identify 150 kinds of element labels and count the proportion of elements (Yao et al. 2019).

According to the research needs of this study, green plant elements (including 5.tree, 10.grass, 18.plant; flora; plant life, 73.palm; Palm tree) is used to calculate the green visibility, and the calculation formula of the GVI of each observation image is:

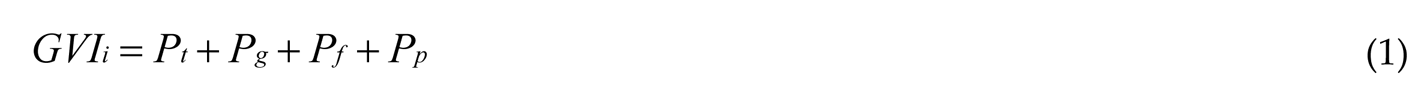

where *GVI_i_* is the GVI in the i^th^ direction, *P_t_* is the proportion of trees counted by the image semantic segmentation software, *P_g_* is the proportion of grassland, *P_f_* is the proportion of flora and plant life, and *P_p_*is the proportion of palm plants.

Python software is used to collect street view photos from 0°, 90°, 180° and 270° directions in BSV Map from a human’s horizontal perspective as GVI observation images. Use visual image semantic segmentation software to analyze the proportion of green plant elements to calculate the GVI_i_ of each observation image, and then average the GVI of the four observation images at each observation point, which is the GVI of this observation point (Figure 5). The calculation method of GVI at each observation point is as follows:

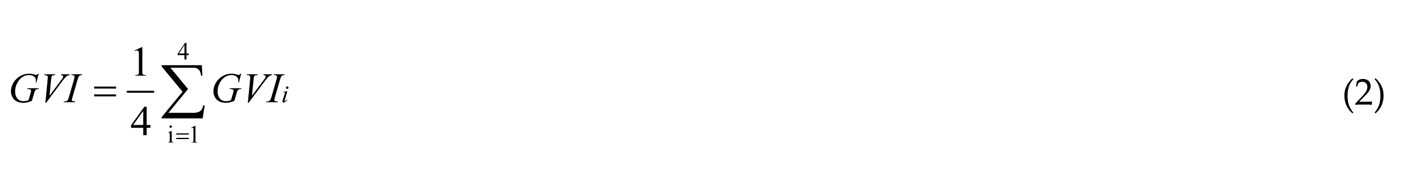

**Fig. 5.**
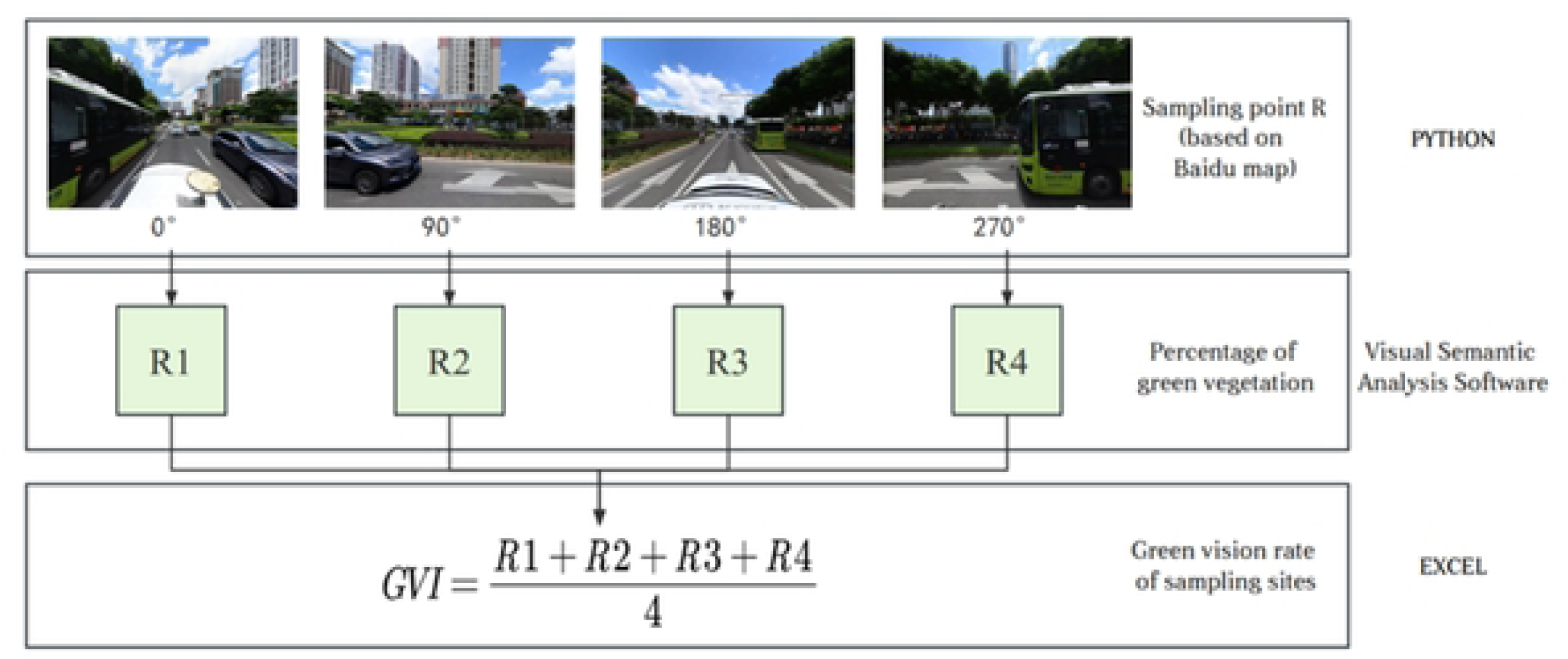
Diagram of calculation method of GVI.

### Normalized difference vegetation index (NDVI)

Normalized difference vegetation index (NDVI) is the most common vegetation remote sensing index that quantifies vegetation by measuring the difference between near infrared (strong vegetation reflection) and red light (vegetation absorption) (Pettorelli et al. 2005). The specific formula is: NDVI=(NIR-R)/(NIR+R), where NIR is the pixel value in the near infrared band, and R is the red band. The indicator range is [-1,1]. In this study, domestic GF-1 satellite images with multispectral resolution better than 8 m were used to calculate NDVI. Based on ENVI 5.3 software, download and install China Satellites in the APP Store to load the GF-1 image, then carry out preprocessing such as radiation calibration and atmospheric correction, use the band math tool to perform band calculation. For the band attributes of the GF-1 satellite, enter the formula (float (B4) - float (B3))/(float (B4)+float (B3)).

### Establishment of observation point buffer and NDVI partition statistics

A large number of studies have confirmed that the occurrence, spatiotemporal distribution, mutual coupling, and other characteristics of landscape pattern processes are scale dependent, and the vegetation index should also have spatial scale characteristics (Bao and Alexis 2022). Therefore, this study selects nine spatial scales, 8m, 16m, 25m, 50m, 100m, 150m, 200m, and 250m, to calculate the NDVI mean value, as a foundation for the correlation study between NDVI and GVI. This study is based on the ArcMap 10.4 to set the buffer for each observation point (Analysis > Proximity > Buffer). The pavement of each road is plotted in the 91 Satellite Map Enterprise Edition (the 19 level image of the 91 Satellite Map can achieve a resolution of 0.28 meters). The boundary of the pavement is not bounded by the actual road width, but is drawn from visual analysis to the impermeable interfaces (walls, buildings, etc.) within a range of 250 meters (Figure 2). Cut the buffer zone from the road surface to obtain the buffer zone for each observation point (Analysis Tools > Extract > Clip), and then perform spatial analysis (Spatial Analysis Tools > Zonal > Zonal Statistics as Table) to export the attribute table.

### Correlation analysis between GVI and NDVI

Research on correlation is the most basic content of science. In today’s era of massive data, research on the correlation between massive data of things has become a hot spot in scientific research (Fan et al. 2014). In previous studies, NDVI and GVI were often used to represent two-dimensional and three-dimensional green indicators respectively, but there was little research on the correlation between them. In this paper, Pearson Correlation Coefficient is used to calculate the correlation between the NDVI mean value and GVI at different scales. Pearson Correlation Coefficient was proposed by Karl Pearson in the 1880s, is widely used to measure the degree of correlation between two variables, and its value is between -1 and 1. The specific formula is (Puth et al. 2014):

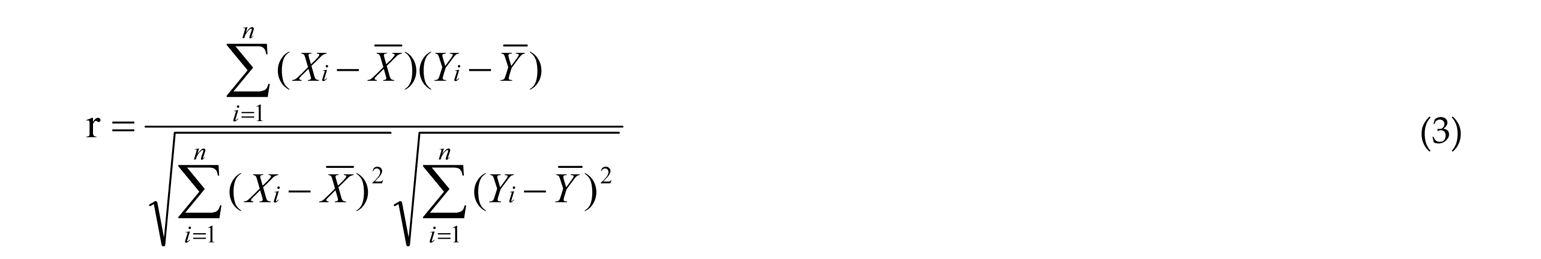

Where r is Pearson Correlation Coefficient, and the closer the value of r is to 1, it means that the two variables are positively correlated, and the stronger the linear correlation is; The closer to -1, the negative correlation; Close to or equal to 0 means that the linear relationship between two variables is weak or not.

## Results

### Statistics of GVI

GVI of each observation point is statistically obtained by using the green visibility calculation method shown in Figure 5. Among all the observation points (Figure 6), the point with the highest GVI is the 8th point of Hongle Avenue in Torch Development Zone Street, which is near the Taxation Bureau and Deneng Lake Park (coordinates are 113.4732443_22.55471916), and the GVI reaches 50.06%. The lowest is at the northern end of Choi Hung Avenue in the West District Street, between the traffic control office and the fire squadron, and the GVI is only 0.26% (coordinates are 113.323077_22.57351983).

**Fig.6.**
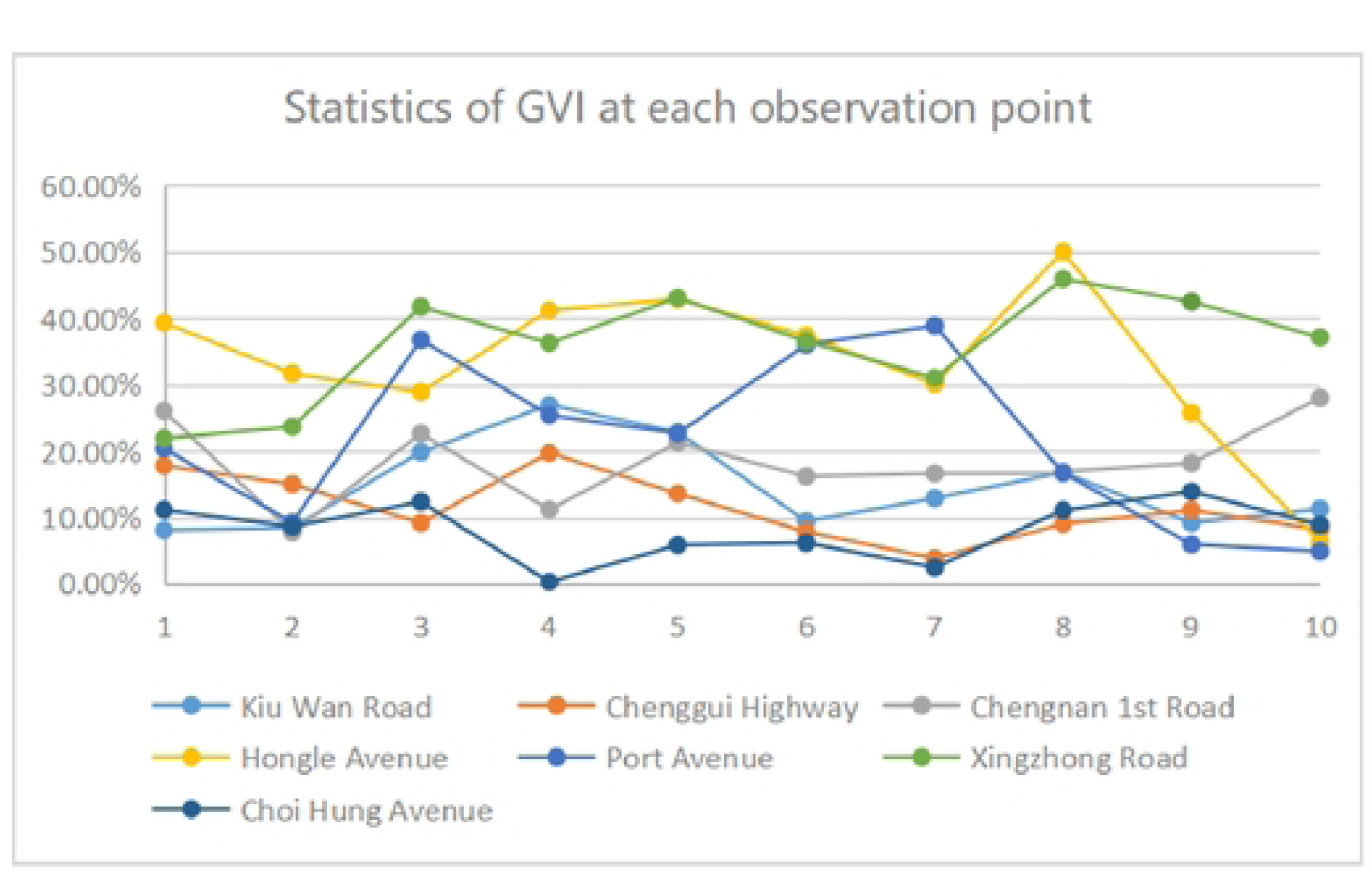
Statistics of GVI at each observation point

The average GVI of street landscape is the highest in Xingzhong Road of Shiqi District, reaching 36.00%, and the lowest in Choi Hung Avenue in West District, only 8.06% (Figure 7). The average GVI of other roads is arranged from high to low, namely, Hongle Avenue in Torch Development Zone is 33.37%, Port Avenue of Port Town is 21.69%, Chengnan 1st Road of South District Street is 18.48%, Kiu Wan Road of East District Street is 14.57%, and Chenggui Highway of Wuguishan District Street is 11.51%. Studies have shown that when the GVI of road greening landscape is more than 30%, such green road space can be welcomed by people (Aoki 1991). It can be observed that the green landscape of Xingzhong Road of Shiqi District Street and Hongle Avenue of Torch Development Zone Street is more comfortable and secure, while the comfort of other streets needs to be improved from the perspective of visible green.

**Fig. 7.**
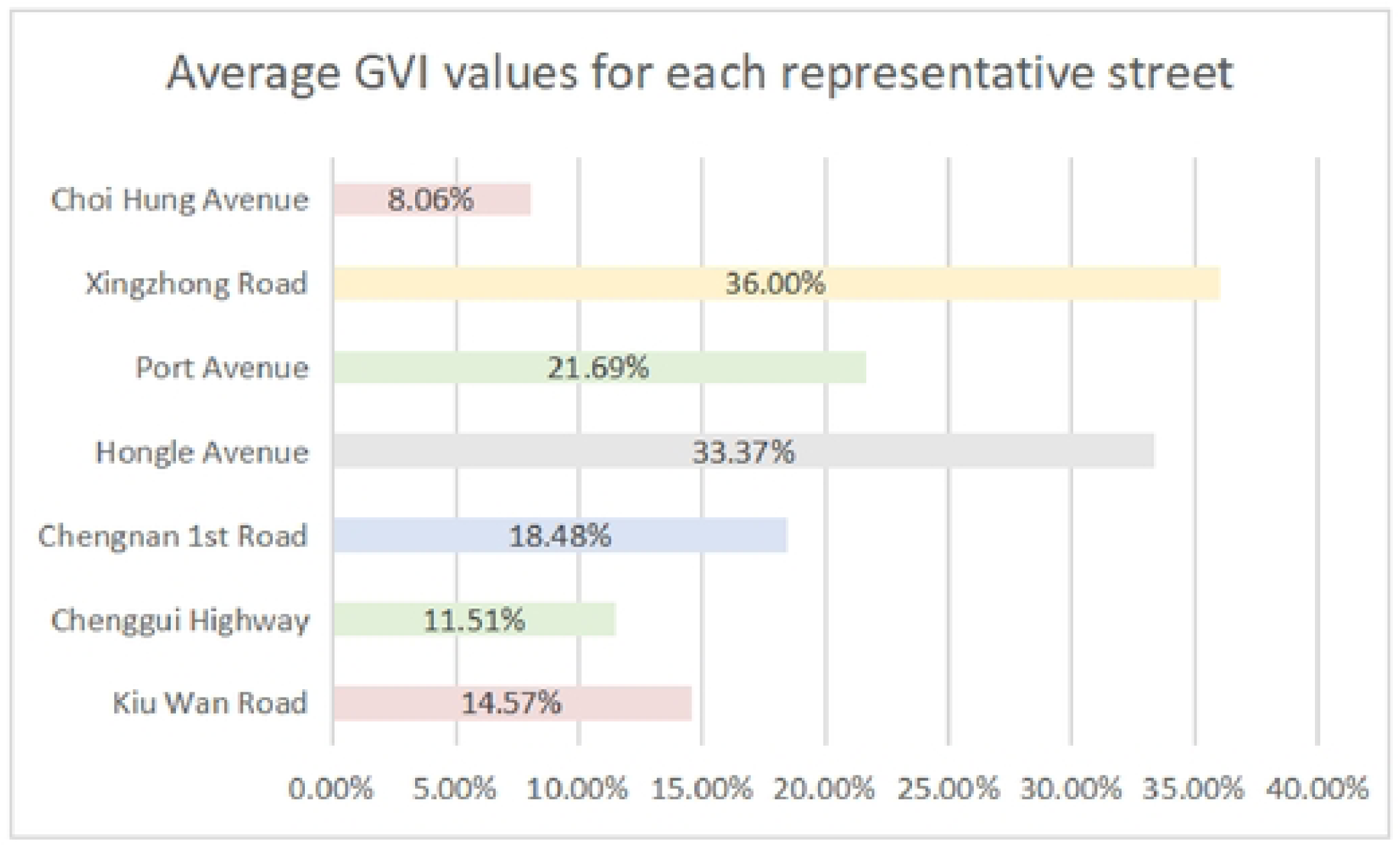
Average GVI values for each representative street.

### NDVI statistics of observation point buffer

Calculate the NDVI mean values of the nine spatial scales of each observation point (8m, 16m, 25m, 50m, 100m, 150m, 200m and 250m) respectively, and draw the box chart (Figure 8) and trend chart (Figure 9) of each spatial scale. It can be found that the NDVI mean values of each street observation point also show different changes with the increase of the buffer scale, showing scale sensitivity. It is mainly divided into three categories: 1) Xingzhong Road, Hongle Avenue, Chengnan 1st Road, and Port Avenue. The NDVI of this kind of roads reaches the peak at 16-50m, and then gradually decreases. For example, when NDVI mean value is 0.42 at the 8m scale of Xingzhong Road, the NDVI mean value is 0.45 at the 16m and 25m scales, and then it shows a downward trend, and NDVI mean value at the 100m to 250m scale does not change significantly. The NDVI mean value is 0.34 at the 8m scale of Hongle Avenue, 0.35 at the 16m scale and 0.34 at the 25m scale, and the downward trend of NDVI is very mild at the 100m to 250m scale. 2) Kiu Wan Road and Choi Hung Avenue. The NDVI of this kind of roads show a trend of decreasing at first and then increasing. The NDVI at 25m scale is the lowest, but the index of Kiu Wan Road changes significantly, while the index of Choi Hung Avenue changes very little. 3) Chenggui Highway. the NDVI of this kind of roads always shows an increasing trend from 8m to 250m. The NDVI of Chenggui Highway in 8m is only 0.09, and in 250m is 0.17.

**Fig. 8.**
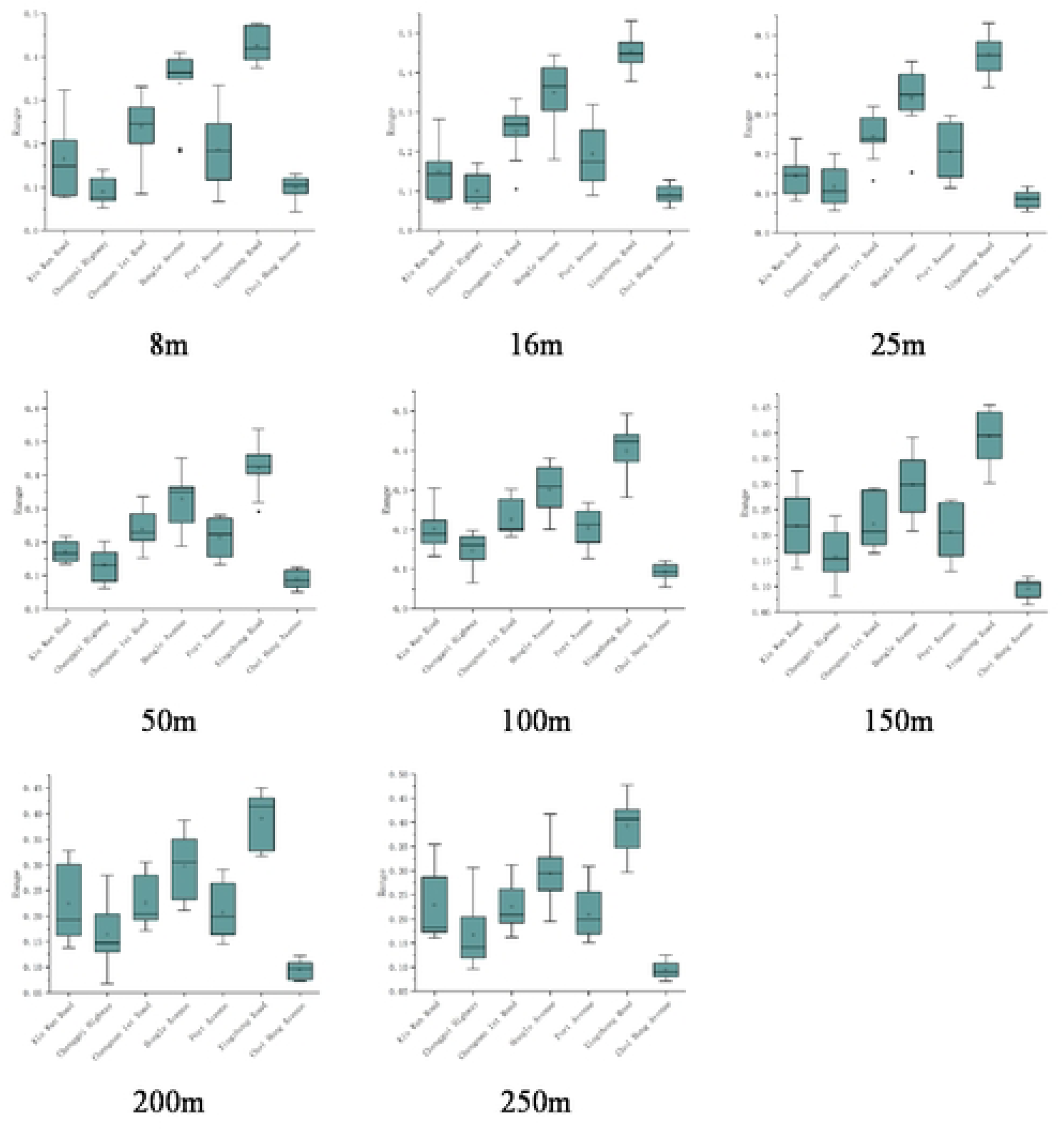
Distribution of NOVI mean values at various observation points within buffer zones of different scales.

**Fig. 9.**
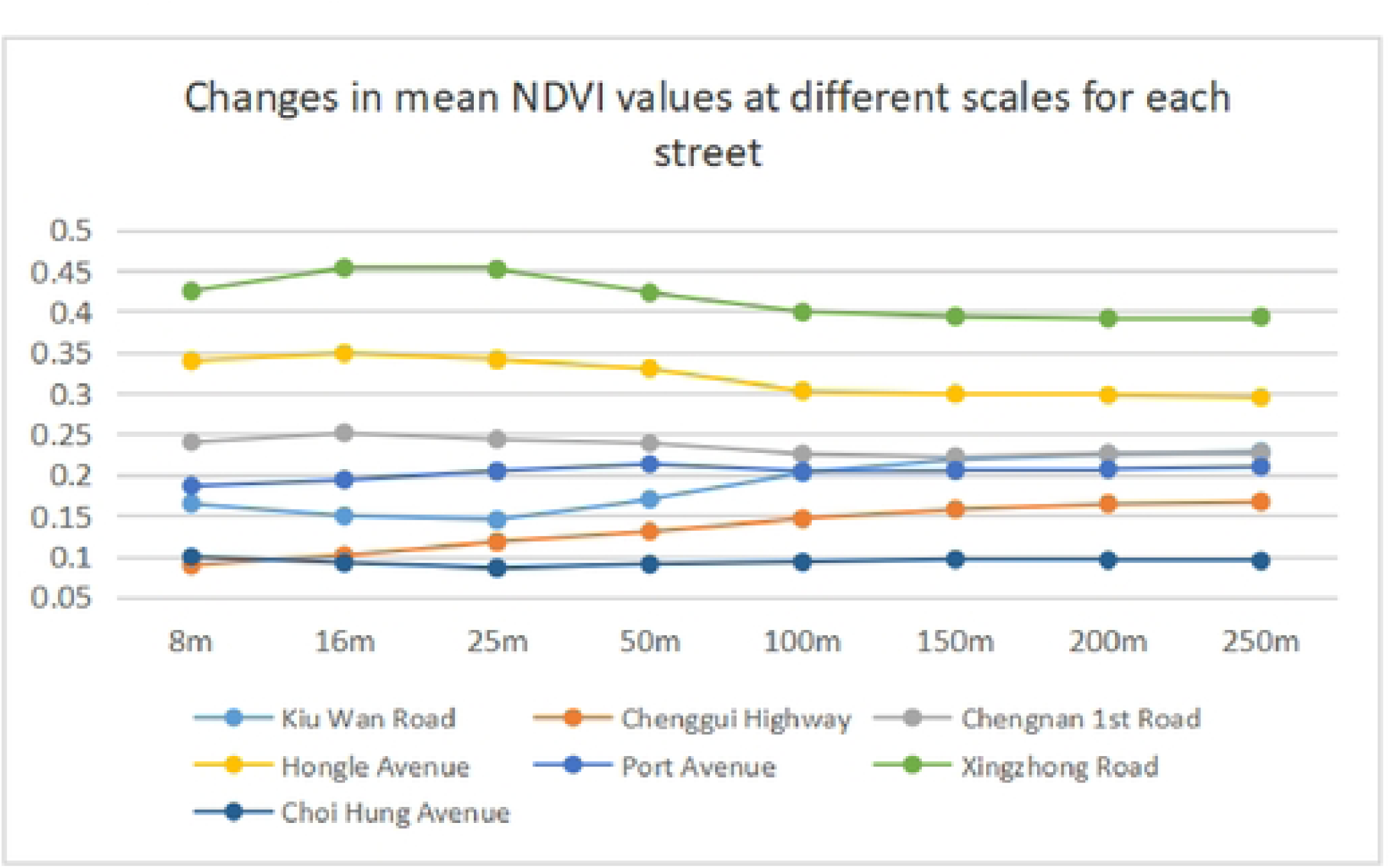
Changes in mean NOVI values al different scales for each street.

Observing the distribution of the NDVI mean value of each street, the ranking is slightly different under each research scale. Taking the 8m scale as an example, the order of the NDVI mean value from big to small is: Xingzhong Road of Shiqi District Street, Hongle Avenue of Torch Development Zone Street, Chengnan 1st Road of South District Street, Port Avenue of Port Town, Kiu Wan Road of East District Street, Chenggui Highway of Wuguishan District Street, and Choi Hung Avenue of West District Street. Judging from the average evaluation results of GVI and NDVI, Xingzhong Road is the most important urban green axis in the downtown area of Zhongshan, and Hongle Avenue has also formed a green street interface of major cities with regional characteristics, which can be used as a typical representative of urban regional streets.

### Correlation analysis between GVI and NDVI

Correlation analysis requires the data to obey normal distribution. Based on origin 2022, this study examines the data distribution of GVI and NDVI mean value of different scales by drawing distribution map and adding whiskers. The results are shown in Figure 10, which all meet normal distribution and can be used for correlation analysis.

**Fig. 10.**
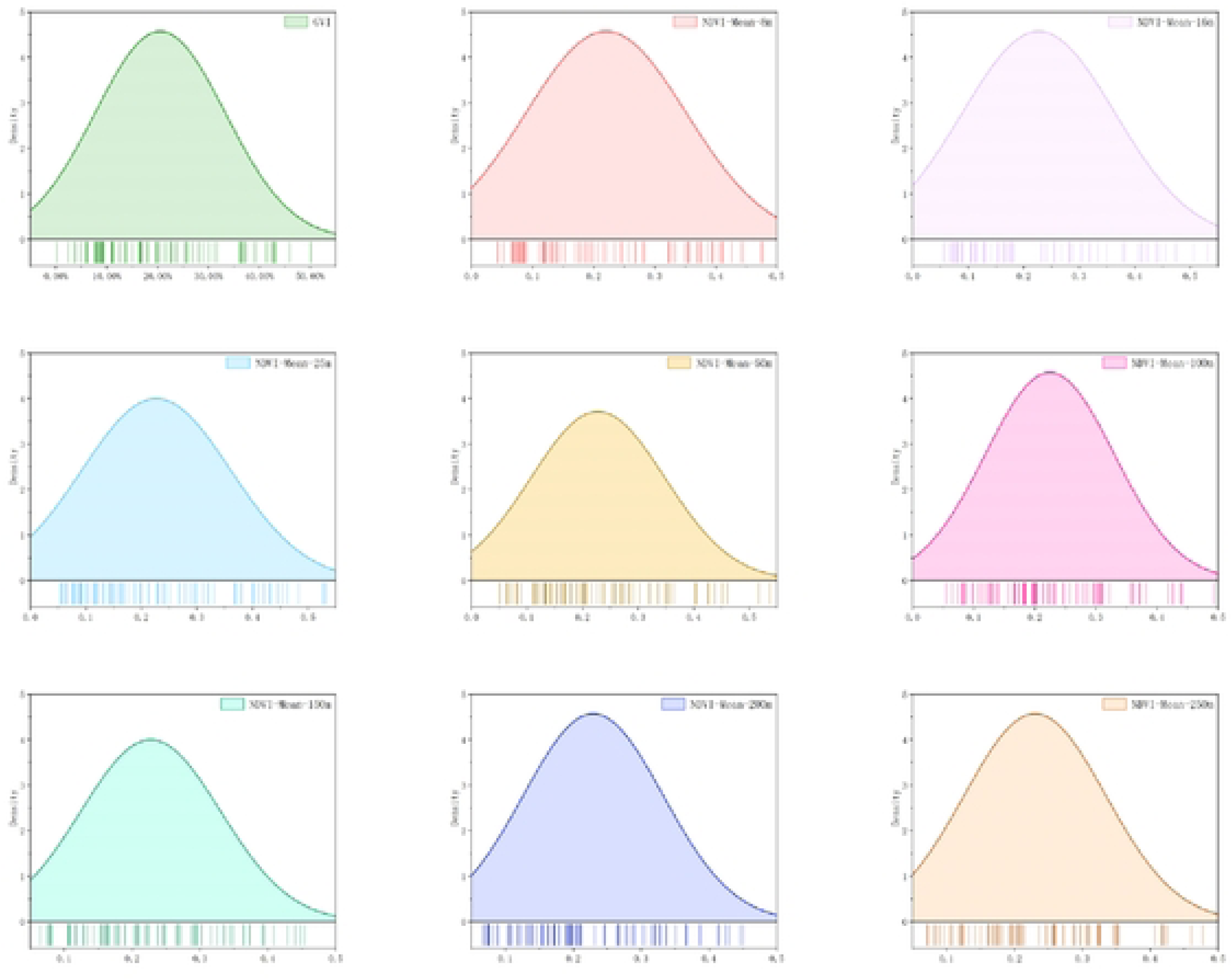
Normal distribution tesl of NOVI values for GVI and different scale buffers.

IBM SPSS Statistics 26 software is used to analyze the Pearson Correlation Coefficient between NDVI mean value and GVI at different scales, specifically: Analyze > Correlate > Bivariate, check Pearson Correlation Coefficient to operate, and the specific results are shown in Table 1. It shown that the NDVI and GVI are highly correlated on the scale of 8m-50m, and Pearson Correlation Coefficient are both greater than 0.8. From 8m to 25m, the correlation showed an increasing trend, and the correlation coefficient reached its peak at 25m, and then showed a decreasing trend. From the experimental results, there is a significant correlation between the street green viewing rate and the average NDVI in Zhongshan City, that is, streets with high NDVI values will also have higher GVI.

**Table 1.** Correlation analysis results.

| Analysis scale (m) | Pearson Correlation Coefficient | Conspicuousness |
| --- | --- | --- |
| 8 | 0.844 | ** |
| 16 | 0.855 | ** |
| 25 | 0.862 | ** |
| 50 | 0.843 | ** |
| 100 | 0.781 | ** |
| 150 | 0.746 | ** |
| 200 | 0.719 | ** |
| 250 | 0.666 | ** |

### GVI prediction model based on NDVI

Based on the above research, the NDVI mean value of streets is highly correlated with the GVI on the scale of 8m-50m, and the correlation coefficient is the highest on the scale of 25m. Therefore, the NDVI of streets on the scale of 25m is selected for linear fitting with GVI, and a GVI prediction model based on NDVI is constructed (Figure 11). For seven representative streets in the central city of Zhongshan, GVI=0.8249*NDVI+0.0181, R^2^=0.7433.

**Figure.11.**
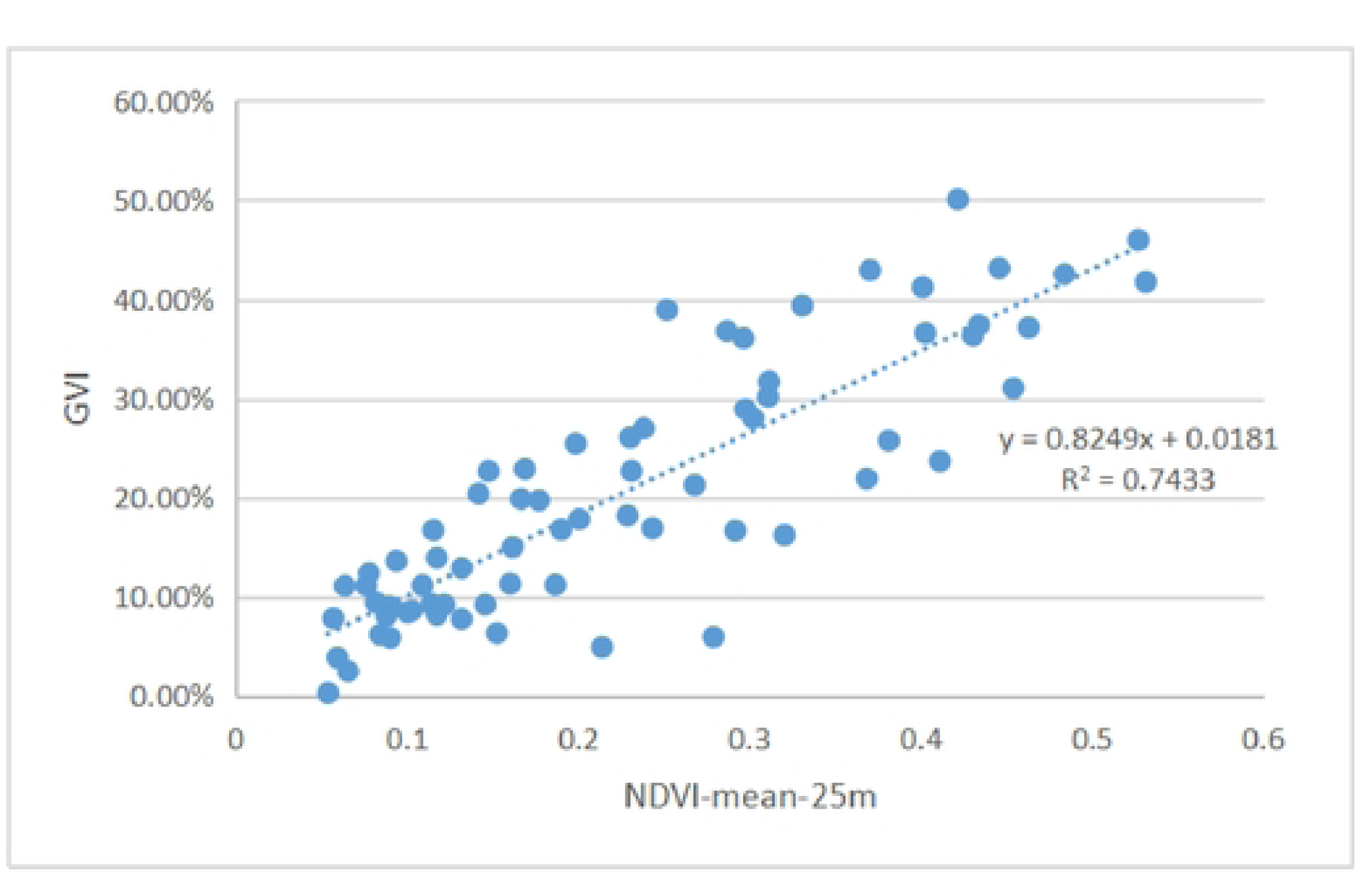
Construction of GVI model based on NDVI

### Analysis of factors affecting the GVI of urban streets

#### The Effect of Street (Town) Functions on GVI

Due to the different tasks and development directions undertaken by each Street (Town) in the development process, and the different greening structures, there are significant differences in the GVI. However, due to the fact that urban administrative divisions do not necessarily follow geographical boundaries, a city is not a divided area, but rather a whole. Many areas and streets blend with each other, and the greening landscape is also sustainable. Therefore, this study only compares the regions with the highest and lowest GVI.

From the mean value of GVI, within the main urban area of Zhongshan, the administrative area has a high GVI. Take Shiqi District Street as an example, which is close to Qijiang River and is one of the central urban areas of Zhongshan. It has unique geographical and development advantages. Within the jurisdiction, supporting services such as administration, commerce, education, medical care, and leisure are complete, and attention is paid to the street landscape and residential environment. The results of GVI to some extent predictable. In contrast, the West District Street mainly serves as an important transportation hub and manufacturing and trade market in Zhongshan. Guangzhu Light Rail, National Highway 105, Guangzhu West Line Expressway, Jiangzhong Highway, Xinqijiang Highway, North Outer Ring Road, Boai Road, Fuhua Road, Zhongshan Road, and other roads cross the region vertically and horizontally. The West District Street has developed commerce and trade, including the earliest developed small commodity market in the Pearl River Delta region, the largest professional computer market in the city, and the wholesale market for agricultural products. Due to the high density of transportation and industrial air pollution, it can have a negative impact on the health of plants and reduce their leaf density, making it more difficult to achieve a high GVI. This area has a low GVI. Generally speaking, the GVI of streets in urban administrative and commercial areas is higher than that in transportation hubs and manufacturing areas.

#### Influence of street width and layout on GVI

Generally speaking, the closer a plant is to the street, the higher its GVI. This is because non green vegetation, such as buildings, roads, and cars, can form visual barriers that block the view of green plants from certain angles or distances. Therefore, when there are no plant barriers between the lanes, the wider the road, the lower the GVI. For example, the width of Choi Hung Avenue along the river section is larger than that of urban streets, and there are fewer plant barriers in this section, so the GVI of Choi Hung Avenue is lower than that of urban streets.

#### Effects of plant species and quantity on GVI

Different plant species have different leaf densities and heights, which can affect the visual effect of greening. For example, tall trees can provide a three-dimensional sense and enhance the three-dimensional effect of urban greening. In contrast, low shrubs or ground cover plants do not have the same visual effect, but can still contribute to the overall greening landscape rate. Secondly, the diversity of plant species is also an important factor affecting the GVI. Using plant species of different heights in green space can create a more dynamic landscape, thereby improving the overall GVI. In addition to planting general broadleaf trees, Xingzhong Road and Hongle Avenue also planted tropical trees such as palm trees. The street has a wide traffic distribution bandwidth, and there are various types of shrubs and ground cover plants, with the highest average GVI on street.

### Deficiencies in the perception of street greening

According to the analysis of the street landscape (Table 2), it can be observed that the greening landscape of some streets in the central urban area of Zhongshan is good, but there are also some street greening landscapes with many problems.

**Table 2.** Street landscape analysis.

| Street | Street type | Greening form | Street tree | Green visual ratio evaluation | NDVI mean (50m) |
| --- | --- | --- | --- | --- | --- |
| Kiu Wan | Two way eight lane | Simple | Trees+Shrubs | Relatively low | Relatively low |
| Port | Two way eight lane | Relatively rich | Trees+Shrubs | Relatively high | Normal |
| Hongle | Two way four lane | Rich | Palm+Trees+Shrubs | High | High |
| Chengnan 1st | Two way six lane | Relatively rich | Trees | Normal | Relatively high |
| Xingzhong | Two way six lane | Rich | Palm+Trees+Shrubs | Highest | Highest |
| Chenggui | Two way eight lane, and some parts are under construction | Simple | Trees+Shrubs | Low | Low |
| Choi Hung | Two way eight lane | Simple | Trees | Lowest | Lowest |

The statistical results of each street landscape show that the GVI in administrative areas in Zhongshan is generally higher than that in industrial areas and transportation hubs. On urban streets (Xingzhong Road), the green volume of the streets is in a dual high state (high GVI and high NDVI index), which is due to the higher level of greening maintenance achieved in areas close to the central urban area. The closer it is to an industrial area (such as Choi Hung Avenue), the negative impact of high-density transportation and industrial air pollution on the plant state will reduce its leaf density, making it more difficult to achieve a high green appearance ratio. The road is also in a dual low state of GVI and NDVI index.

Currently, the focus of improving the environmental quality of streets in Zhongshan is on the maintenance and widening of roads, while the scientific conservation of plants is lacking. Some green belts were not immediately restored after removal to facilitate construction, or were not further maintained after restoration. Taking Chenggui Highway as an example, Wugui Mountain Street is different from other streets. As the only town street in the whole city that has been classified as an ecological protection zone, it has the only contiguous mountain system in the core area of the Greater Bay Area of Guangdong, Hong Kong, and Macao. The mountain forest area is 88 square kilometers, and the forest coverage rate reaches 81.2%. It is known as the “Green Lung of Zhongshan”. However, among randomly selected streets, their GVI is only higher than that of Choi Hung Avenue in the West District Street. The main reason is that due to road construction, the greening area has been greatly reduced, resulting in aging and out-of-grade plants.

In the study, it was also found that in road landscapes, the GVI of streets with multiple plants selected for matching is generally higher than that of streets with single plant species. Xingzhong Road and Hongle Avenue are rich in plant configurations, including trees, shrubs of different colors and shapes, and ground cover plants. However, there are still many sections in Zhongshan where the plant configuration is relatively monotonous and lacks characteristics in order to consider economic issues and ensure survival rates.

## Discussion

This study uses BSV Map and semantic segmentation to study the perception of urban street green from the perspective of combining GVI and NDVI, and conducts a correlation study between GVI and NDVI from different scales, which is helpful for comprehensively and scientifically evaluating urban street landscape greening and providing optimization suggestions. Compared to similar studies (Li et al. 2021; Toikka et al. 2020), this study has certain advantages in using deep learning software to analyze GVI: 1) visual image semantic segmentation software no longer requires tedious manual operations, but directly obtains results through software, so there is almost no error calculation. 2) because the recognition of plants does not entirely rely on color features, the plant color or other colored plants affected by the shooting time will also be accurately calculated. However, this method is not perfect, and is mainly subject to interference from two aspects: data sources and plant species. In terms of data sources, the problem with the python algorithm used in this study is that it can only obtain images with a fixed resolution size. A clearer photo resolution often leads to better recognition results. Semantic segmentation of visual images is not a software specifically designed to calculate the GVI, and there are only a few plant elements in it. Whether or not it can maintain accurate recognition rates in the face of more complex plant species has not yet been tested. However, this situation is rare, and these four elements are completely capable of completing most tasks.

Landscape patterns are scale dependent, therefore, scale and scale change are key to understanding and studying landscape patterns. As a remote sensing vegetation index, NDVI also have scale effects, so selecting the scale at which to obtain NDVI values becomes the key to this study. Compared to similar studies (Li et al. 2021; Lv et al. 2022), which mostly use Landsat 8 OLI remote sensing images with a resolution of 30 meters as the data source for NDVI calculation, this study has greater advantages in selecting domestic GF-1 satellite images with a multispectral resolution better than 8 meters for studying street landscape greening. This study selects nine spatial scales, namely, 8m, 16m, 25m, 50m, 100m, 150m, 200m, and 250m, for analysis. The upper limit is selected as 250m, mainly considering that the maximum distance that the human eye can clearly see the contour of an object is 250m, which is to ensure the maximum consistency with the GVI observation. On larger scales (50m to 250m), use 50m as the scale interval. Within 50m, 8m, 16m, and 25m are selected as the analysis scales. The minimum of 8m is because the multispectral resolution of the GF-1 satellite image selected for the project is 8m, and 16m is multiplied by a 2-fold relationship based on 8m. The 25m scale is the distance that people can just see each other’s faces in external space, which is also the unit modulus that can be used in external space research mentioned by Yoshinobu Ashihara in his “External Modulus Theory”. In addition, there are also some differences between this study and other researches in terms of buffer zone delimitation. For example, Lv Xuxin (2022) used the width of the road to add 50m buffer zone to both sides to calculate the NDVI mean value of each street, and then analyzed the correlation and influencing factors between the GVI and NDVI. Due to the differences in the nature of land used on both sides of different streets, some roads within a 50m buffer zone may have types of street green space and institutions green space, while others within a 50m buffer zone may already have types of residential buildings or commercial buildings. This delimitation of the latter clearly has certain drawbacks, as people’s sight cannot penetrate the buildings, however, it is obviously not scientific to include buildings and green spaces behind buildings in the statistics of buffer zones as a study of the correlation with GVI. Therefore, when setting the research scope in this study, it is not a one-size-fits-all approach. Instead, based on the actual situation on both sides of the street, the research scope of each road is drawn using artificial visual judgment in the 91 Satellite Map Enterprise Edition (0.28 meter resolution). The research scope is not bounded by the actual street width, but is drawn from visual analysis to the impermeable interfaces (walls, buildings, etc.) within 250 meters, NDVI statistics are calculated within the range obtained after cutting the buffer zone and road extent. Such a range setting helps to study the relationship between NDVI and GVI in a more detailed manner, but there are also some shortcomings. Due to the heavy workload of the road boundary delimitation based on artificial vision judgment and rendering in this topic, the sample data is relatively limited.

This study indicated that the NDVI mean value of street observation points varies with the increase of buffer size, and NDVI is scale sensitive. Most streets have a significant inflection point in the NDVI mean value at the scale of 16-25m, and the NDVI mean value at the scale of 100-250m has a relatively flat change trend. Only the Chenggui Highway is a special case, and the NDVI mean value of this street has always shown an increasing trend between the scale of 8-250m. To analyze the reasons, in order to obtain higher accuracy in the experimental method set up in this study, when delimiting the observation range, as long as obstacles such as surrounding buildings or walls are encountered, the subsequent vegetation coverage rate will be ignored (because it cannot be seen, it is not related to the GVI). There are many buildings on both sides of the streets in the central urban area, and the vegetation behind the buildings and buildings is not included in the statistics. Therefore, the change trend of NDVI mean values on the 100-250m scale of most streets is basically relatively flat, and the change mainly comes from the NDVI changes in the front and rear directions of the street, rather than the left and right directions. Chenggui Highway is located in Wuguishan District Street. Due to the establishment of a national nature reserve in Wuguishan, the surrounding buildings of the town’s streets are relatively limited, and people’s sight can also reach a far range. Therefore, the NDVI mean value in Wuguishan has always shown an increasing trend.

At the same time, this study calculated the Pearson Correlation Coefficient between the NDVI mean value and the GVI at different scales, and found that the NDVI mean value and the GVI are highly correlated on the 8-50m scale, and the correlation coefficient at the 25m scale is the highest, reaching 0.862. Based on this, this study constructed a model that can quickly predict GVI, which is of practical significance in compensating for the lack of street view maps in some regions or time periods and thus unable to quickly calculate the GVI. And provide new ideas for urban environmental assessment, spatial optimization, and landscape improvement. Because street view maps are not perfect in terms of spatial coverage and temporal variability (Kim and Jang 2023), for example, it cannot be guaranteed that street view maps are available at every period, and there is insufficient spatial coverage of street view images for some pedestrian based commuter routes. Therefore, it is very important to study the correlation between NDVI and GVI. In the future, the relationship between street NDVI and GVI should be further refined and in-depth studied, such as exploring the scale sensitivity of urban street landscape NDVI to GVI by expanding the research scope, increasing sample selection, and conducting classification and grading discussions on streets. For different road types and street levels, the best scale for research should be selected, and GVI prediction models for various street scenes should be constructed respectively. So as to better provide scientific support for the optimization of urban street landscape pattern.

In order to improve the quality of street greening on a humanistic scale, based on the above analysis of the characteristics of GVI and NDVI, and the shortcomings in the perception of street green volume, the author proposes the following optimization strategies: 1) Strengthen the foresight of green space system planning, and promote the integrated design of both inside and outside the red line of the street. Based on the analysis results of GVI and NDVI, identify the streets that need to be focused on, clarify the width of green lines on both sides of the street in the green space system planning, and promote the integrated design of roadside green belts and ancillary green spaces on surrounding plots, thereby enriching the level of green landscape and improving the level of GVI. 2) Timely remedy plant “breakpoints”. After the repair is completed, the damaged street plants should be restored as soon as possible to increase the continuity of green. Regularly prune older trees in the streets to ensure that plants have reasonable growth space, improve species diversity, and stabilize the ecological environment. The streets of Zhongshan have distinctive characteristics, with numerous streets and lanes crisscrossing each other. Obviously, many branch streets can no longer be planted with large plants such as trees. At this time, can rely on climbing plants to cover the street walls, expand three-dimensional greening, and increase the GVI; Or by designing a pocket park and landscape sketches to connect the green areas into one. 3) Change the planting strategy. Taking the West District Street of Zhongshan as an example, increasing GVI does not necessarily require rare tree species, but rather can select plants with strong vitality for the purposes of cooling, blocking dust, reducing noise, and purifying the air. The temperature in Zhongshan is generally high, and the midday sun is strong. A higher GVI can significantly alleviate visual fatigue, block strong light, reduce acoustic and light pollution, and improve the driving safety of transportation hubs. At the same time, it can improve the stereotype of citizens and tourists on manufacturing areas.

## Conclusions

As a city with complex street structures and a long history of construction, the planning of Zhongshan is more complex than that of other cities. Based on the deep learning full convolutional network visual image semantic segmentation software trained with ADE_20K dataset, this study provides a simple, convenient and fast data source that can quickly evaluate street green perception and analyze street landscape in a short period of time. Exploring street green perception from the perspective of combining GVI and NDVI is helpful for scientifically evaluating the greening effect of urban street landscapes, identifying existing problems, and providing optimization plans. Studying the correlation between GVI and NDVI, and selecting appropriate scales to build models for rapid prediction of GVI, can help solve the shortcomings of street view maps in terms of spatial coverage and temporal variability. Although this study still has some limitations, such as that representative streets are mostly two-way eight lanes, and the number of sample points selected is limited, the idea of this article is very important for studying the relationship between urban street GVI and NDVI. In particular, selecting an appropriate research scale and constructing a GVI prediction model through NDVI not only provides a rapid method for estimating GVI, but also solves the practical difficulties of missing street view maps for some streets and parts of the time, which has practical significance. This study provides new ideas for urban environmental assessment, spatial optimization, and landscape improvement.

